# Distal-less genes *Dlx1/Dlx2* repress oligodendrocyte genesis through transcriptional inhibition of *Olig2* expression in the developing vertebrate forebrain

**DOI:** 10.1101/2020.04.09.012385

**Authors:** Qiang Jiang, Jamie Zagozewski, Roseline Godbout, David D. Eisenstat

## Abstract

In this study, we demonstrate that mouse DLX2 binds to the *Olig2* gene locus in mouse embryonic forebrain *in vivo*. We further confirm the specificity and the transcriptional repressive effect of the binding *in vitro*. Furthermore, loss of *Dlx1/2* function leads to increased *Olig2* expression in the ventral embryonic mouse forebrain *in vivo*. As well, we demonstrate that chicken DLX1 binds to one chicken Olig2 gene domain *in vitro*, and overexpression of *Dlx1* is sufficient to repress *Olig2* expression in the developing chicken forebrain *in ovo*. Chicken DLX1 with a mutation eliminating its DNA binding ability is unable to bind to the *Olig2* probe *in vitro*, and abrogates its repressive function on *Olig2* expression *in ovo*. Our results establish that *Dlx1/2* is both necessary and sufficient to repress oligodendrocyte specification mediated via direct transcriptional inhibition of *Olig2* expression in the developing vertebrate forebrain.

## INTRODUCTION

Both neurons and the principal types of glia-oligodendrocytes and astrocytes-in the vertebrate central nervous system (CNS) are derived from the same progenitor pool. One fundamental question in developmental neurobiology is how the vast diversity of neuronal and glial subtypes is generated from the relatively simple and undifferentiated neuroepithelium of the embryo (Rowitch & Kriegstein, 2010). It has been established in mouse that the same progenitor domains within the ventral telencephalon give rise to both cortical inhibitory interneurons and oligodendrocytes (Rowitch & Kriegstein, 2010; Wonders & Anderson, 2006). The ventral telencephalon is therefore viewed as a suitable model system to address the factors that determine neuron-glial fate decisions, and further the mechanisms controlling the balance between neurogenesis and oligodendrogenesis during CNS development (Petryniak, Potter, Rowitch, & Rubenstein, 2007).

Most, if not all, inhibitory cortical interneurons in rodents arise from the developing ventral forebrain and migrate tangentially to reach their cortical destination, although the precise origin of interneuron populations in human remains controversial because of the evidence that dorsal progenitors may produce a proportion of the cortical interneurons (Hansen et al., 2013; Letinic, Zoncu, & Rakic, 2002; Molnar & Butt, 2013; Yu & Zecevic, 2011). The vertebrate *Dlx* genes, a family of homeodomain containing transcription factors, are essential regulators of the normal migration and differentiation of progenitors of the subcortical telencephalon (Panganiban & Rubenstein, 2002). In the mouse, four of the *Dlx* genes, *Dlx1, Dlx2, Dlx5* and *Dlx6* are expressed in an overlapping and sequential pattern in the developing forebrain (Eisenstat et al., 1999; Liu, Ghattas, Liu, Chen, & Rubenstein, 1997). *Dlx1/Dlx2* double mutants exhibit a severe block in neurogenesis in the subcortical telencephalon (Anderson, Qiu, et al., 1997; Marin, Anderson, & Rubenstein, 2000), a massive reduction of the inhibitory interneurons of the cerebral cortex, hippocampus and olfactory bulb due to defects in tangential cell migration (Anderson, Eisenstat, Shi, & Rubenstein, 1997; Long et al., 2007; Pleasure et al., 2000), as well as accelerated and increased oligodendrogenesis in the ventral embryonic forebrain (Petryniak et al., 2007).

In vertebrates, it is now evident that both region-restricted and temporal cell-intrinsic regulators operate to segment the neuroepithelium into progenitor domains for sequential specification of distinct neuron and glial cell types. Generally, gliogenesis follows neurogenesis with switching of developmental programs in the same progenitor domains (Rowitch & Kriegstein, 2010). This was first observed in the ventral spinal cord, where lines of evidence indicated that oligodendrocyte production is initiated in the same pMN region that generates motor neurons. The basic helix-loop-helix (bHLH) transcription factors encoded by the genes *Olig1* and *Olig2* further provide the molecular mechanisms that connect the development of motor neurons and oligodendrocytes (Rowitch, 2004). Deletion of both *Olig1* and *Olig2* resulted in significant loss of motor neurons in the spinal cord and total elimination of oligodendrocyte formation throughout the CNS (Q. Zhou & Anderson, 2002). Furthermore, a single gene mutation of *Olig2* can lead to a similar failure of pMN domain development and specification of motor neurons and oligodendrocytes in the spinal cord (Lu et al., 2002; Takebayashi et al., 2002).

Oligodendrocytes within the mouse telencephalon are generated in three spatiotemporal waves. In the mouse, the first oligodendrocyte precursor cells (OPCs) are derived from the medial ganglionic eminence (MGE) and anterior entopeduncular area (AEP) at embryonic day 11.5 (E11.5), followed by a second wave from the lateral and/or caudal ganglionic eminences (LGE, CGE) starting at E15.5, and finally a third wave from the cortex after birth (Kessaris et al., 2006). During forebrain development, the OPCs are coupled to GABAergic interneurons, rather than motor neurons. Clonal analysis indicated that the basal forebrain contains common progenitors capable of producing both interneurons and OPCs (He, Ingraham, Rising, Goderie, & Temple, 2001; Yung et al., 2002). In addition, basal forebrain derived *Dlx* expressing cells can give rise to oligodendrocytes in the dorsal telencephalon (He et al., 2001; Marshall and Goldman, 2002). Furthermore, Petryniak et al reported that *Dlx1/2* could repress OPC formation by negatively regulating *Olig2* expression, and determine GABAergic interneuron versus oligodendrocyte specification from common progenitors in the ventral embryonic forebrain, although *Dlx* expression does not represent an irreversible state of neuronal commitment (Petryniak et al., 2007). However, whether the effect by DLX1/2 on *Olig2* expression is mediated directly or indirectly during forebrain development remains to be determined.

In this study, we demonstrate the first report of *in vivo* binding of DLX2 protein to the *Olig2* gene locus in mouse embryonic forebrain using chromatin immunoprecipitation (ChIP). We further confirm the specificity and the transcriptional repression effect of the binding *in vitro*. Moreover, loss of *Dlx1/2* function leads to increased Olig2 expression in the ventral forebrain of mouse embryos. We also show that chicken DLX1 (cDLX1) binds to one chicken *Olig2* gene domain *in vitro*, and overexpression of *cDlx1 in ovo* is sufficient to repress Olig2 expression in the developing chicken forebrain. In addition, mutation of chicken DLX1 resulting in elimination of its DNA binding ability is unable to bind to *Olig2 in vitro*, abrogating its ability to repress *Olig2* expression *in ovo*.

## MATERIALS AND METHODS

### Animals and tissue preparation

All animal protocols were conducted in accordance with guidelines set by the Canadian Council on Animal Care, and approved by the Animal Care and Use Committee of the University of Alberta. For mouse experiments, embryonic age was determined by the day of appearance of the vaginal plug (E0.5), confirmed by morphological criteria. Wild type tissues were collected from CD-1 mice (Charles River Laboratories), and mice with null mutations of *Dlx1* and *Dlx2* are a kind gift from Dr. Rubenstein (University of California San Francisco)(Qiu et al., 1997). For chicken experiments, fertilized White Leghorn chicken eggs were obtained from the poultry unit of the University of Alberta. Tissues were fixed with 4% paraformaldehyde (PFA) in phosphate buffered saline (PBS, pH 7.4) at 4°C overnight, infiltrated in 20% sucrose, and embedded with VWR Clear Frozen Section Compound. Coronal sections at 12 µm were prepared for further applications.

### Cloning of chicken *Dlx* genes

Chicken *Dlx1* was cloned by means of RT-PCR, with the poly A^+^ RNA from E7 chicken retina used as the template. The cDNA was generated through a 3’-RACE-Ready cDNA synthesis reaction using the SMARTer RACE cDNA amplification kit (Clontech), following the user manual. The PCR primers are forward: **5’-GCACACACACAGACCCAGCGGCGAG-3’**, reverse: **5’-CGTGTTCGGGCTCACATAAGCTGCGG-3’**. The PCR product was ligated into the pGEM-T Easy vector and verified by direct sequencing. For chicken *Dlx2*, two pairs of PCR primers were designed according to the predicted sequence in the updated chicken genome assembly (Gallus_gallus-5.0) to amplify the potential chicken *Dlx2* with the same cDNA template used for cloning chicken *Dlx1*, forward 1: **5’-ATGACCGGCGTCCTGCCGAGCC-3’**, reverse 1: **5’-CTAGAAGATGCCGCCGCCGCCG-3’**, forward 2: **5’-CCTCGGGATGACCGGCGTCCTG-3’**, reverse 2: **5’-GTCTTCCGCAAAGGCACCTATCGAC-3’**.

### *In ovo* electroporation

The open reading frame (ORF) of chicken *Dlx1* was subcloned into the adapter plasmid SLAX 12 NCO at the *NcoI* and *EcoRI* sites, and then further into the *ClaI* site of the avian replication-competent retrovirus vector RCASBP(B) (Li, Monckton, & Godbout, 2014) with the In-Fusion HD cloning kit (Clontech). The mutant construct that converts glutamine (Q) to glutamic acid (E), by altering the DNA codon CAG to GAG at amino acid position 50 (Q50E) of the chicken DLX1 homeodomain (amino acid 177 of the chicken DLX1 protein) was also generated using the In-Fusion HD cloning kit with the same vector. *In ovo* electroporation was carried out at E2 (Hamburger Hamilton stage 10-11) using equipment described by Li (Li et al., 2014). Briefly, RCASBP(B), RCASBP(B)-Dlx1, or RCASBP(B)-Dlx1 Q50E plasmids were injected into the lumen of each embryo’s forebrain vesicle at a concentration of 2 µg/µl with Fast Green. Electrodes were placed at 4 mm apart on either side of the embryonic forebrain, and five 20 volt, 50 msec square pulses were applied unilaterally. Electroporated embryos were further incubated for 3 days, and harvested on E5 after green fluorescent protein (GFP) expression screening.

### Chromatin immunoprecipitation (ChIP) assay

ChIP assays were carried out as described (Q. P. Zhou et al., 2004) with minor modifications. Briefly, tissue of E13.5 mouse ganglionic eminences (GEs) and hindbrain (does not express *Dlx* genes) were harvested and crosslinked using 1% PFA with protease inhibitors for 1.5 hour at room temperature (RT). Cross-linked cells were resuspended in lysis buffer (50 mM HEPES pH 7.5, 140 mM NaCl, 1 mM EDTA pH 8.0, 1% Triton X-100, 0.1% Sodium Deoxycholate, 0.1% SDS with protease inhibitors) and sonicated. After 8000 g centrifugation for 5 minutes at 4°C, Pierce Protein A/G UltraLink Resin (ThermoFisher Scientific, 53132) was added to the supernatant to pre-clear the chromatin. Repurified DLX2 antibody (Eisenstat et al., 1999; Porteus et al., 1994) was then added to the chromatin solution and incubated at 4°C overnight. On the following day, Pierce Protein A/G UltraLink Resin was added and incubated for 4 hours at 4°C. Beads were then pelleted and sequentially washed with Low Salt Wash Buffer, High Salt Wash Buffer twice, LiCl Buffer, and TE buffer twice. The enriched DLX2 DNA complexes were eluted and reversal of the cross-links was performed at 65°C overnight. Finally, proteinase K was added for 2 hours incubation at 65°C, and DNA was purified using Qiagen PCR purification kit (Cat No. 28106). The ChIP enriched genomic DNA was then used as the template to perform PCR amplification, with oligonucleotide primer pairs targeting seven regions of the mouse *Olig2* sequence (ENSMUSG00000039830) upstream of its translation start site. PCR products were purified after gel electrophoresis, and then ligated into the pGEM-T Easy vector (Promega) for sequencing verification.

### Electrophoretic mobility shift assay (EMSA)

Potential regions of the mouse *Olig2* locus as DLX2 binding targets were screened by ChIP PCR assays, and cloned into the pGL3-Basic vector. The restriction fragments of each region were then gel purified and labeled with [α^32^P]-dGTP (PerkinElmer) using the large fragment of DNA polymerase I (Invitrogen). For further analysis of the putative TAAT/ATTA homeodomain binding motifs within the target region, synthetic complementary oligonucleotides were obtained from Integrated DNA Technologies (IDT, Edmonton, Canada), annealed and then labeled with [γ^32^P]-ATP (PerkinElmer) in the presence of T4 Polynucleotide Kinase (Invitrogen). Radioactively labeled probes were purified using MicroSpin G-25 columns (GE Healthcare). The binding reaction mixture was prepared in a 20 μl volume, containing 80,000 cpm labeled probe, 1 µg poly (dI-dC), and 200 ng purified recombinant mouse DLX2 protein in Gel Shift Binding Buffer (Promega). Unlabeled probe was added for ‘cold’ competition, while the specific DLX2 antibody was added for ‘supershift’ assays. The DNA-protein complexes were resolved on a 4% non-denaturing polyacrylamide gel after incubation.

For chicken EMSA assay, the pET-21b vector was used as the expression construct for chicken DLX1 and DLX1 Q50E proteins. After transformation, BL21 competent cells were cultured in MagicMedia expression medium (Invitrogen) for protein production. Both chicken DLX1 and DLX1 Q50E proteins were purified using Capturem His-Tagged Purification Maxiprep Kit (Clontech), and verified by Western blot with primary antibodies against 6x His tag (Invitrogen, MA1-21315), and DLX1 (Abcam, ab167575). Synthetic oligonucleotide probes were prepared as described above. In a 20 μl volume, 400 ng purified chicken DLX1 or DLX1 Q50E protein was added. Other binding and electrophoresis conditions were the same as for mouse EMSA experiments.

### Reporter gene assay

An effector plasmid expressing mouse DLX2 was constructed as described (Q. P. Zhou et al., 2004). Reporter plasmids were constructed by inserting the mouse *Olig2* gene mR2 and mR6 regions identified by the ChIP assays *in situ* into the pGL3-Basic vector (Promega), respectively. Correctly constructed plasmids were confirmed by direct DNA sequencing. Transient co-transfection experiments were performed in the HEK 293 human embryonic kidney cell line (courtesy of Dr S. Gibson, University of Manitoba). Cells at 90 % confluence in 24-well plates were transfected with 0.5 μg effector plasmid, 0.5 μg luciferase reporter plasmid, together with 0.1 μg pRSV-β-gal plasmid (Promega) as a control for transfection efficiency, using Lipofectamine 2000 (Invitrogen). Cells were subsequently harvested 48 hours later. Luciferase activity was measured with the Luciferase Assay System (Promega, E1501) and a standard luminometer, and normalized with β-galactosidase activity.

### *In situ* hybridization (ISH)

For ISH, the chicken *Dlx1* containing pGEM-T Easy plasmids were linearized, and riboprobes were prepared using the DIG RNA labeling mix and SP6/T7 transcription kit (Roche). Sections were fixed in 4% PFA for 10 minutes, washed in dimethyl dicarbonate (DMPC, Sigma) treated PBS. Slides were then placed in freshly prepared acetylation solution: 0.75 ml triethanolamine (Sigma) and 125 µl acetic anhydride (Fisher Scientific Canada) in 50 ml H_2_O/DMPC for 10 minutes, and washed with PBS/DMPC. The riboprobes were diluted in hybridization solution: 50% formamide (EMD Millipore), 5x Saline Sodium Citrate (SSC) solution, 10% dextran sulfate (EMD Millipore), 1x Denhardt’s solution (Invitrogen), 250 µg/ml yeast tRNA (ThermoFisher Scientific) and 0.1% Tween 20 (TW) in H_2_O/DMPC, preheated and applied to sections. Hybridization was performed at 65°C overnight. The post-hybridization washes were done 3 times in preheated wash buffer: 50% formamide, 1x SSC solution, 0.1% TW at 65°C, each time 15 minutes. After cooling to room temperature, slides were rinsed in maleic acid buffer containing TW (MABT, pH 7.5), blocked in blocking buffer with 2% Blocking Reagent (Roche), 20% goat serum in MABT, and then incubated in 1:1500 dilution of anti-digoxigenin-AP, Fab fragments (Roche) in blocking buffer. Color development was achieved using NBT/BCIP (Roche) in alkaline phosphatase buffer (NTMT).

For combination of ISH with immunohistochemistry (IHC), rabbit anti OLIG2 (EMD Millipore, AB9610) 1:200 was added together with the anti-digoxigenin-AP, Fab fragments after routine ISH was started as described above. After an overnight incubation, the IHC was first completed using EXPOSE rabbit specific HRP/DAB detection IHC kit (Abcam, ab80437). Sections were then washed and further subjected to color development for ISH.

### Immunofluorescence (IF) assay

IF experiments were performed as previously described (de Melo, Qiu, Du, Cristante, & Eisenstat, 2003) with slight modifications. Primary antibodies used were: rabbit anti-DLX2 (Eisenstat et al., 1999; Porteus et al., 1994) 1:300, rabbit anti-OLIG2 (EMD Millipore, AB9610) 1:200, goat anti-OLIG2 (Santa Cruz Biotechnology, sc-19969) 1:500. All Alexa Fluor secondary antibodies (Invitrogen) were used at 1:200: donkey anti-rabbit 488/594, donkey anti-goat 594. Sections were cover-slipped using VECTASHIELD mounting medium with DAPI (Vector laboratories), and examined with a Nikon Eclipse TE2000-U inverted microscope.

### Reverse transcription-quantitative PCR (RT-qPCR) assay

The tissue of E13.5 mouse GEs was dissected, and the total RNA was extracted using TRIzol (Invitrogen). To synthesize complementary DNA (cDNA) via RT, SuperScript III reverse transcriptase (Invitrogen) was used with 500 ng RNA as the template, and oligo (dT) as the primer. qPCR was performed on a LightCycler 96 System (Roche), using the FastStart Essential DNA Green Master (Roche), with 100 ng cDNA and 1 μM of each primer in a 20 μl reaction volume. PCR conditions were: pre-incubation, 95°C 600 s; amplification, 95°C 10 s, 55°C 10 s, and 72°C 10 s, for 45 cycles; melting, 95°C 10 s, 65°C 60 s, and 97°C 1 s. The qPCR experiments were accomplished on three technical replicates of three matched littermate pairs of wild type (WT) and *Dlx1/2* double knockout (DKO) mouse embryos. The difference of the expression level of *Olig2* mRNA between WT and DKO embryos was evaluated by means of comparative quantification of the quantification cycle (Cq) values, using mouse *Gapdh* gene as the internal reaction control. The Student’s t-test was used to assess the statistical significance of the difference.

## RESULTS

### *Dlx1/2* and OLIG2 are co-expressed in a subset of progenitors within the basal embryonic telencephalon

In chicken, we focused our studies on *cDlx1* due to lack of *Dlx2* sequence information. We first investigated expression of *cDlx1* RNA in coronal sections at rostral and intermediate telencephalon levels of E5 chicken embryos by ISH. The chicken *Dlx1* probe strongly labels the subventricular zone (SVZ) of the subpallial striatum (ST), and pallidum (PA) domains. *cDlx1* can also be faintly detected in the ventricular zone (VZ) and mantle zone (MZ) of both ST and PA (**Figure 1**, A, C, E). We further examined the expression of chicken OLIG2 protein by IF analysis in adjacent E5 intermediate forebrain sections. In contrast to *cDlx1*, OLIG2 is primarily expressed in the PA domain, and restricted to nearly all cells of the VZ, as well as part of the SVZ (**Figure 1**, D, E). In order to determine whether *cDlx1* and OLIG2 are co-expressed in chicken, we combined ISH with IHC on intermediate forebrain sections of E5 chicken embryos. Robust co-localization of both *cDlx1* ISH and OLIG2 IHC signals was detected in the SVZ of the PA domain, at the boundary of OLIG2 expression, with double staining also present in scattered cells in the VZ of the same PA domain (**Figure 1**, E, E’).

**Figure 1.**
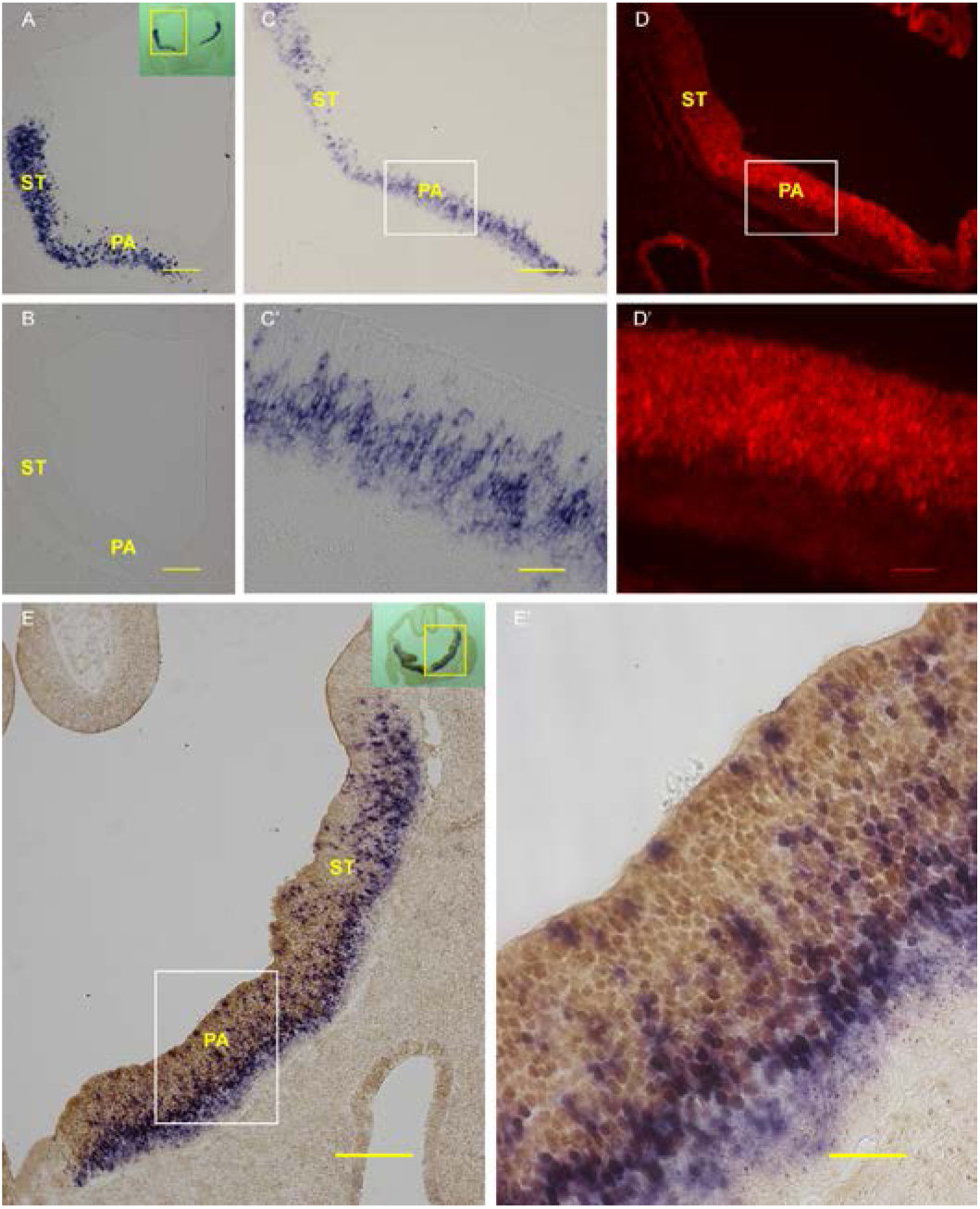
Co-expression of *cDlx1* and OLIG2 in the ventral embryonic chicken forebrain. **A-C**, *cDlx1* expressing cells labeled by ISH in ST and PA domains in coronal sections through rostral level (**A**), with rostral section incubated with sense probe as a negative control (**B**), and intermediate level of E5 chicken forebrain (**C**); **C’**, is the boxed PA area in (**C**) at higher magnification, showing the predominant expression of *cDlx1* in the SVZ. **D**, OLIG2 expression visualized using IF in the PA region in E5 chicken intermediate forebrain section directly adjacent to (**C**); **D’**, is the higher-magnification view of the boxed region in (**D**), demonstrating the restriction of OLIG2 expression in the VZ, as well as part of the SVZ. **E**, co-expression of *cDlx1* and OLIG2 on coronal E5 chicken intermediate forebrain section, visualized as overlap of *cDlx1* ISH signal (blue) with OLIG2 IHC signal (brown) in the SVZ and VZ of the PA region; **E’**, is the image at higher magnification of the (**E**) boxed region. Insets in (**A**) and (**E**), lower magnification view of the same whole section, respectively, with the box representing the position of the corresponding image. PA, pallidum; ST, striatum. Scale bars: **A-E**, 100 μm; **C’-E’**, 25 μm.

In mouse, expression of DLX2 protein was closely examined and compared with the expression of OLIG2 protein by double immunofluorescence assay in the developing mouse forebrain. At E13.5, DLX2 expression is well established in the subpallial telencephalon, including LGE, MGE and the septum area at a rostral level (**Figure 2A**, a). Caudally, where the LGE and MGE begin to fuse into the caudal ganglionic eminence (CGE), DLX2 is expressed in the LGE, MGE as well as the anterior entopeduncular (AEP) and preoptic area (POA) (**Figure 2B**, a). In all these anatomic domains (LGE, MGE, septum or AEP/POA), DLX2 is expressed predominantly in the SVZ, with the highest expression detected in the SVZ region adjacent to the VZ/SVZ boundary. DLX2 is also expressed at lower levels in the VZ and the mantle zone (MZ). In addition, DLX2 expression is more abundant in the MZ of the MGE than in the LGE, which is more noticeable on the caudal section (**Figure 2A**, B, a). In contrast, OLIG2 expression is mainly restricted to the MGE rostrally, while at a more caudal level, OLIG2 is expressed in the MGE and the AEP/POA area. At this developmental stage, OLIG2 is intensely expressed in nearly all the progenitors within the VZ. However, there is a dramatic decrease in OLIG2 expression observed in the SVZ, close to the most robust region of DLX2 immunostaining; and furthermore, only sporadic cells are OLIG2 positive in the MZ of both the MGE and AEP/POA (**Figure 2A**, B, b). As a result, cells with co-expression of DLX2 and OLIG2 are mostly localized in the VZ in addition to the adjacent SVZ of the MGE, where OLIG2 appears to decline coincidentally with the observed increase of DLX2 expression. Progressively, in the rest of the SVZ and the MZ, while most of the cells are DLX2 positive, very few cells show double staining of DLX2 and OLIG2 (**Figure 2A**, B, c).

**Figure 2.**
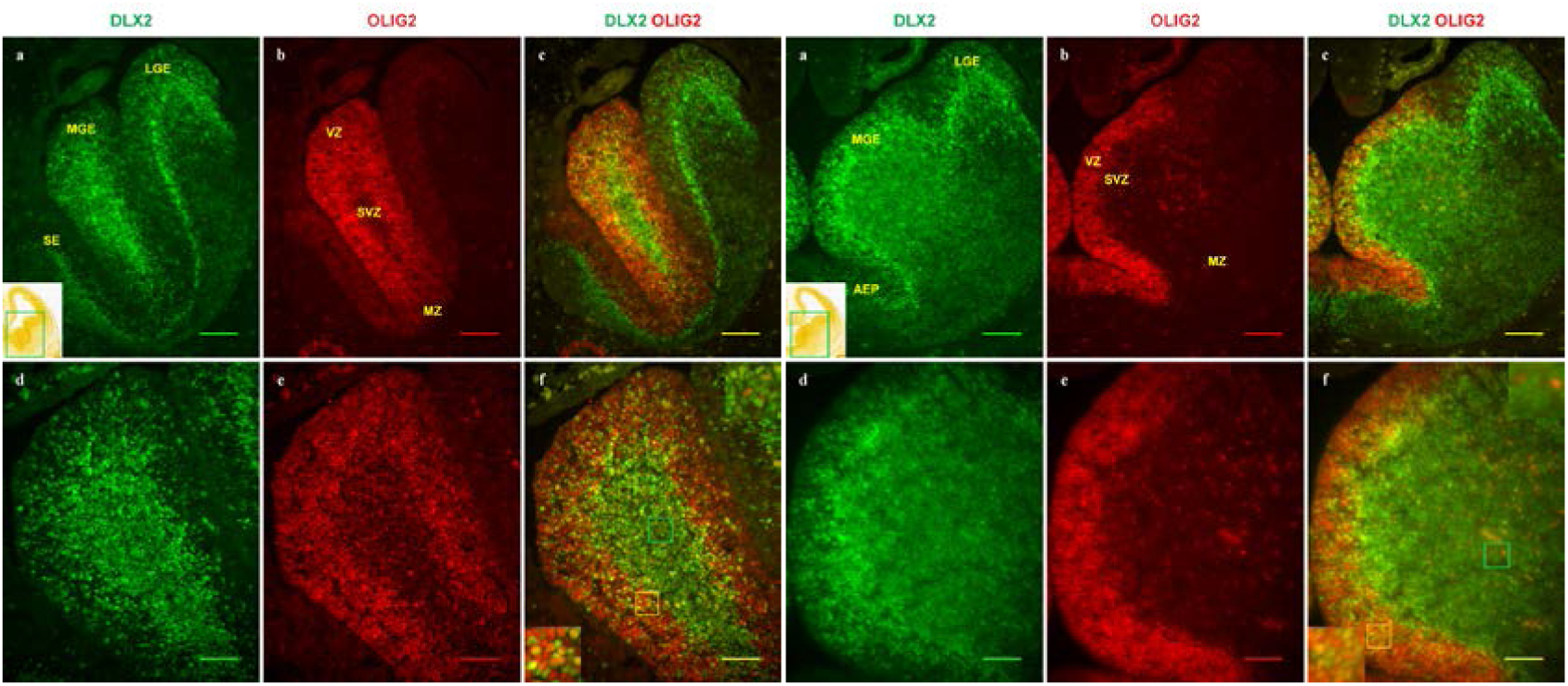
Transient co-expression of DLX2 and OLIG2 proteins in the developing mouse forebrain. **A and B**, Visualization of DLX2 (green), OLIG2 (red) expressing cells in coronal forebrain sections of E13.5 mouse embryos, and cells with co-expression of DLX2/OLIG2 (yellow) in the merged images by double immunofluorescence staining on one rostral section (**A**), and one section at a more caudal level (**B**). The highest expression of DLX2 was detected in the SVZ of LGE, MGE as well as SE (**A, a**) or AEP/POA (**B, a**), while DLX2 is also expressed in the VZ and MZ of all these subpallial domains at a lower level (**A, B, a**). At this stage, OLIG2 is primarily restricted to the VZ of the MGE at a rostral level (**A, b**), and the VZ of the MGE and AEP/POA more caudally (**B, b**). In higher magnification views of the MGE (**A, B, d-f**), most cells with co-expression of DLX2 and OLIG2 localize in the VZ and the adjacent SVZ, whereas cells with DLX2/OLIG2 double labeling are rare in the remainder of the SVZ and the MZ (**A, B, f**). Insets in (**A, a**) and (**B, a**) are adapted from the Allen Brain Atlas, indicating position of the images with green boxes; in (**A, f**) and (**B, f**), top right insets show the magnifications of the SVZ regions within the green boxes, while the bottom left insets represent the magnified VZ regions inside the yellow boxes, respectively. AEP, anterior entopeduncular area; LGE, lateral ganglionic eminence; MGE, medial ganglionic eminence; MZ, mantle zone; SE, septum; SVZ, subventricular zone; VZ, ventricular zone. Scale bars: **A, B, a-c**, 100 μm; **d-f**, 50 μm.

Together, our studies in both embryonic chicken and mouse forebrain indicate that *Dlx1/2* and *Olig2* genes are temporarily co-expressed in a common group of progenitors within the ventral embryonic forebrain; however, they tend to be expressed reciprocally as cells exit the progenitor zone and begin to differentiate. These results support that negative regulatory interactions may exist between *Dlx1/2* and *Olig2* during the telencephalon development.

### DLX2 binds to the *Olig2* gene locus in mouse embryonic forebrain *in vivo*

In order to determine whether DLX2 protein directly binds to genomic DNA sequences within the *Olig2* gene locus, ChIP assays with a specific DLX2 antibody was performed on E13.5 mouse GE tissue to isolate the target genomic DNA sequences of DLX2. We then carried out PCR to amplify the predicted DLX2-binding regions in the mouse *Olig2* locus, with oligonucleotide primers flanking seven candidate regions (termed mR1-mR7) upstream of its translation start site based on the presence of putative TAAT/ATTA homeodomain binding motifs (**Figure 3A**). Mouse genomic DNA was utilized as a positive control for the PCR reaction. Omission of the DLX2 antibody served as a negative control for the immunoprecipitation, while E13.5 mouse hindbrain tissue, which does not express any *Dlx* genes, served as a negative tissue control. ChIP-PCR assays yielded a positive signal in regions mR1, mR2, mR5, mR6 and mR7 supporting either direct or indirect promoter occupancy *in vivo*. For all these regions, control assays performed without DLX2 antibody or on *Dlx* negative embryonic hindbrain tissue were negative. The binding specificity of DLX2 to mR2 and mR6 was subsequently confirmed by EMSA assay *in vitro*; thus our ChIP assay findings validated the direct binding of DLX2 to the *Olig2* locus in E13.5 mouse GE tissue (**Figure 3C**). Regions mR1, mR5, and mR7 did not exhibit binding with DLX2 protein in the EMSA assays *in vitro*. We postulate that some DNA fragments after sonication are sufficiently long to overlap the binding region and adjacent non-binding region, and may produce false positive results in the ChIP assay. Data for mR1, mR5, and mR7 are not shown, and these regions were not further characterized. The resulting PCR amplicons were cloned and verified by direct sequencing.

**Figure 3.**
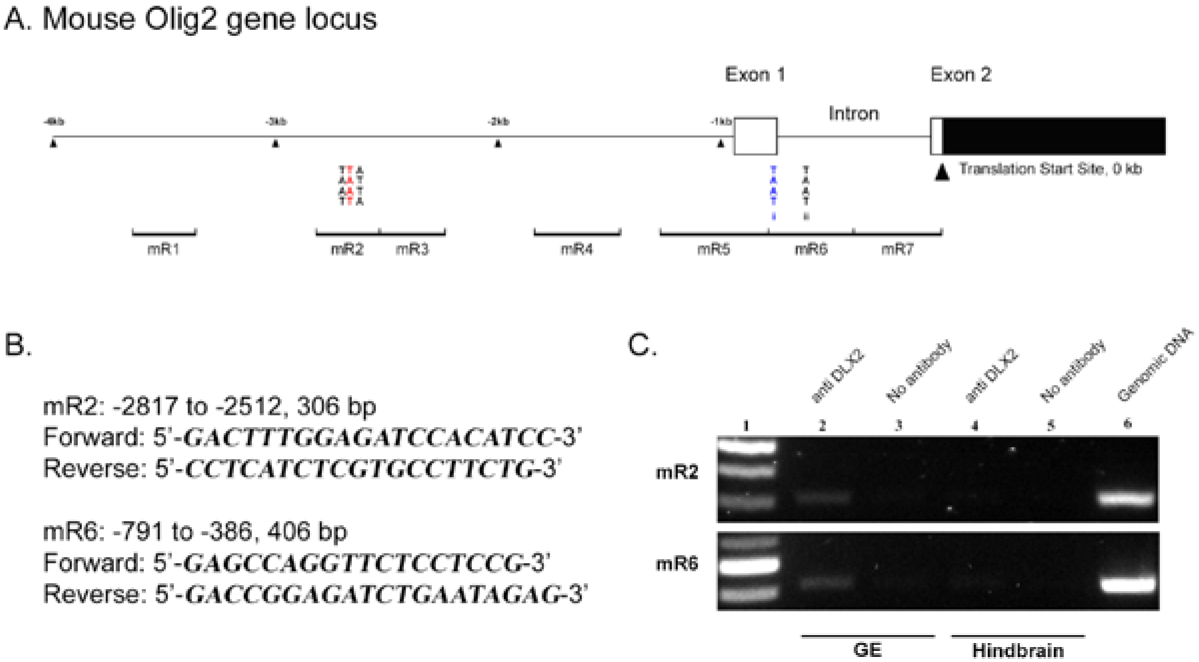
*In vivo* binding of DLX2 protein to the *Olig2* locus in E13.5 mouse forebrain. **A**, Schematic representation of the mouse *Olig2* gene locus. Seven candidate binding regions (designated mR1-mR7), and the putative TAAT/ATTA homeodomain DNA binding sites in mR2 and mR6 are indicated, with respect to the translation start site. **B**, Position relative to the translation start site, the expected size, and the sequence of oligonucleotide primers used for PCR are listed for mR2 and mR6 of the mouse *Olig2* gene. **C**, After isolation of the genomic DNA fragments bound to DLX2 homeoprotein in E13.5 mouse GE tissue using ChIP assays, specific PCR bands were obtained for mR2 and mR6 of mouse Olig2 gene (lane 2). No obvious bands observed in negative controls including ChIP without the addition of DLX2 antibody (lane 3), or ChIP on E13.5 mouse hindbrain, tissue that does not express *Dlx* genes (lanes 4, 5). Mouse genomic DNA was positive as the positive control for PCR reaction (lane 6). GE, ganglionic eminence.

### Chicken DLX1, and mouse DLX2 bind to regions of Olig2 *in vitro*

In order to confirm the binding specificity of mouse DLX2 protein to *Olig2* regions identified by ChIP experiments *in vivo*, we performed EMSAs with recombinant mouse DLX2 protein and radiolabeled probes designed to those corresponding regions. Recombinant DLX2 protein binds to mR2 and mR6 probes, producing bands of less mobile complexes (**Figure 4A**, lanes 2, 6), compared with the probe only controls (**Figure 4A**, lanes 1, 5). The presence of more than one band may be the result of homodimerization of the recombinant DLX2 protein, and/or possible protein modifications. Specific unlabeled probes competitively blocked the binding of the labeled probes, and eliminated shifted bands (**Figure 4A**, lanes 3, *7*). Moreover, the addition of specific anti-DLX2 antibody to the protein-DNA complexes further reduced their mobility and produced supershifted bands (**Figure 4A**, lanes 4, 8). Other candidate homeoprotein binding regions in the mouse *Olig2* locus, mR1, mR5 and mR7, were not significantly bound by DLX2 protein (data not shown). Within the mouse *Olig2* gene mR6, there are two TAAT/ATTA motifs (**Figure 3A**, **Figure 4B**, a). We further identified that mR6i is the specific binding motif of DLX2 protein (**Figure 4B**, b, lane 2). Unlabeled probe competitively inhibited the binding (**Figure 4B**, b, lane 3), and specific DLX2 antibody resulted in the supershift (**Figure 4B**, b, lane 4). Mouse DLX2 protein did not bind to the mR6ii motif of the *Olig2* gene (**Figure 4B**, c).

**Figure 4.**
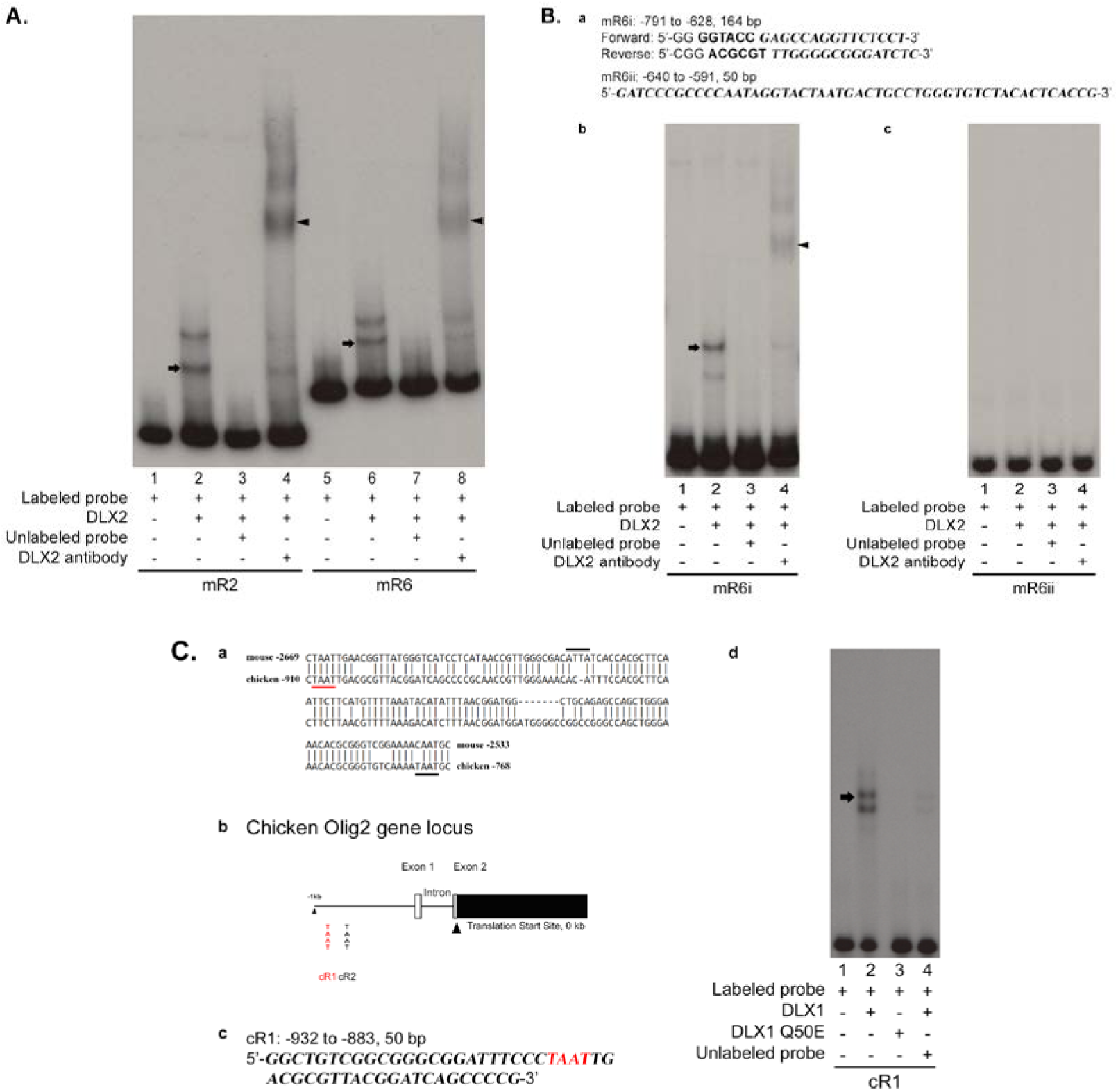
EMSA demonstrated specific binding of DLX1/2 to regions of the *Olig2* gene *in vitro*. **A**, Recombinant mouse DLX2 protein binds to mR2 and mR6 of the mouse Olig2 gene. Radiolabeled probes were incubated alone in lane 1 for mR2, and lane 5 for mR6, respectively; with DLX2 protein in lanes 2-4 (mR2), and lanes 6-8 (mR6). Unlabeled competitive probes were added in lane 3 for mR2, and lane 7 for mR6. Specific DLX2 antibody was added in lane 4 (mR2) and lane 8 (mR6). **B**, Recombinant mouse DLX2 protein binds to the mR6i TAAT motif of the mouse *Olig2* gene. **a**, Position relative to the translation start site, the expected size, and the sequence of oligonucleotide primers used for the cloning of mouse *Olig2* mR6i motif, and the sequence for mR6ii motif oligonucleotide synthesis. The sequence of the mouse Olig2 gene is shown in italics. **b**, Recombinant mouse DLX2 protein binds to the mR6i motif of the mouse *Olig2* gene. Radiolabeled probe was incubated alone in lane 1 and with DLX2 protein in lanes 2-4; with further addition of unlabeled probe in lane 3, or DLX2 antibody in lane 4. **c**, Mouse DLX2 protein does not bind to the *Olig2* mR6ii motif. Lane 1: radiolabeled probe only; lanes 2-4: with DLX2 protein; with further addition of unlabeled probe in lane 3, or DLX2 antibody in lane 4. **C**, Recombinant chicken DLX1 protein binds to the cR1 TAAT motif of the chicken *Olig2* gene. **a**, Conserved region in alignment of the 3 kb genomic sequence upstream of the translation start site of the *Olig2* gene of mouse and chicken. The position of this region relative to the translation start site is labeled. The identical TAAT motif underlined in red corresponds to the second TAAT motif of mR2 in mouse (**Figure 3A**, in red), and cR1 in chicken. This region also contains another potential ATTA motif in mouse (overlined), and one more TAAT motif in chicken only (cR2, underlined). **b**, Schematic representation of the chicken *Olig2* gene locus. Two candidate TAAT binding motifs (cR1, cR2) are indicated with respect to the translation start site. **c**, Position relative to the translation start site, and the sequence for cR1 motif oligonucleotide synthesis. **d**, Recombinant chicken DLX1 protein binds to the cR1 motif of the chicken *Olig2* gene. Lane 1: radiolabeled probe only; lane 2: with DLX1 protein; lane 3: with DLX1 Q50E mutant protein; and lane 4: with DLX1 protein and unlabeled competitive probe. Binding of DLX1 or DLX2 protein to a specific DNA sequence results in gel shifted bands, indicated by arrows (**A**; **B, b**; **C, d**). Binding of DLX2 protein to DNA probe, as well as specific DLX2 antibody produces supershifted bands, indicated by arrowheads (**A**; **B, b**).

Next, we sought to determine the binding site of chicken DLX1 protein in the chicken *Olig2* gene locus. Based on our ChIP and EMSA data in mouse, we aligned 3 kb of genomic sequence immediately upstream from the translation start site of the *Olig2* gene of mouse and chicken, which spans over the mR2 in mouse (**Figure 3A**). The sequence alignment identified one 144 bp conserved region with 76% sequence identity (**Figure 4C**, a). Notably, this conserved sequence resides in part of the mR2 region of the mouse *Olig2* gene, with one identical TAAT motif present in both mouse and chicken (labeled in red, **Figure 3A**, **Figure 4C**, a, b). The two TAAT motifs within the conserved sequence of the chicken *Olig2* gene were named cR1 and cR2. No other TAAT/ATTA motifs were found between cR1 and the translation start site of the chicken *Olig2* gene (**Figure 4C**, a, b). Recombinant proteins of chicken DLX1 and DLX1 Q50E were prepared and verified by Western blot using anti-His tag and anti-DLX1 antibodies (data not shown). Chicken DLX1 protein binds to the cR1 probe of the chicken *Olig2* gene, showing shifted bands of protein-DNA complexes (**Figure 4C**, d, lane 2), whereas DLX1 Q50E mutant protein failed to produce such bands due to loss of its DNA binding ability (**Figure 4C**, d, lane 3) (Le et al., 2007). Again, specific unlabeled probe eliminated the shifted bands by competing with the radiolabeled probe for cDLX1 binding (**Figure 4C**, d, lane 4). Chicken DLX1 protein does not bind to the cR2 region of Olig2 gene (data not shown).

In summary, our EMSA results confirm the binding of mouse DLX2 to mR2 and mR6i regions of the mouse *Olig2* gene *in vitro*. Furthermore, in chicken, DLX1 binds to the cR1 TAAT motif of the chicken *Olig2* gene, which is identical to one conserved TAAT motif in the mR2 region of the mouse *Olig2* gene.

### DLX2 inhibits luciferase reporter gene expression via mR6 *in vitro*

In order to determine the functional significance of DLX2 protein binding to the mR2 and mR6 regions of *Olig2*, we performed luciferase reporter gene assays. Co-transfection of mouse *Dlx2* expressing plasmid resulted in higher luciferase activity of the reporter plasmid with mR2 inserted upstream of the luciferase reporter gene, although without statistical significance (**Figure 5**). However, the luciferase activity of the reporter plasmid containing the mR6 region was significantly reduced when co-transfected with the *Dlx2* construct. These results confirm the functional role of mouse DLX2 as a transcriptional repressor of *Olig2* gene expression via binding at the mR6 domain *in vitro*.

**Figure 5.**
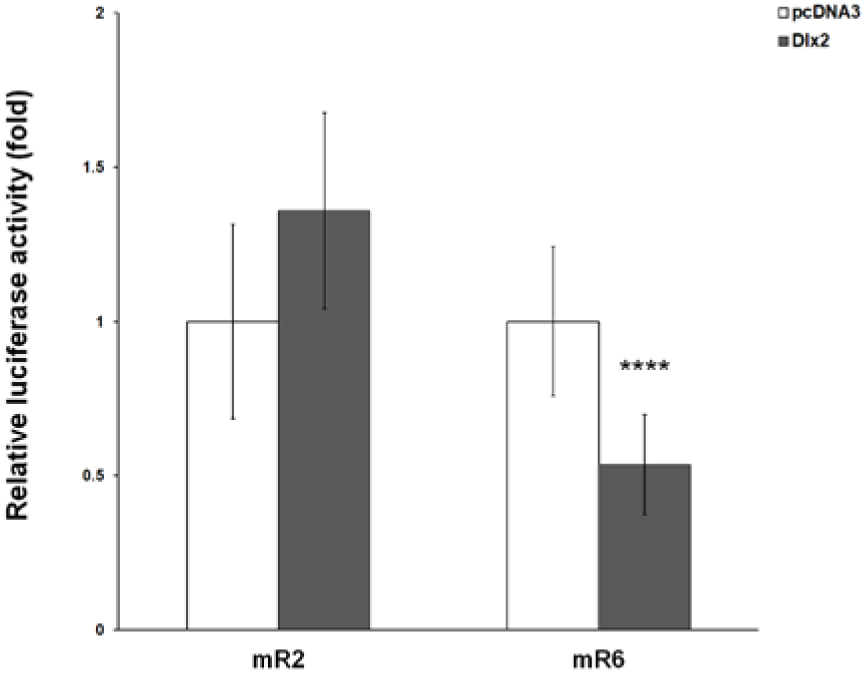
Mouse DLX2 represses transcription of the *Olig2* mR6 reporter construct *in vitro*. In HEK 293 cells, luciferase reporter gene vectors containing either the mR2 or mR6 regions of the mouse *Olig2* gene were co-transfected with an expression construct of *Dlx2*, respectively. *Dlx2* co-transfection leads to significant decrease of luciferase activity for the mR6, but not for the mR2 reporter plasmid. Results are presented as the relative luciferase activity of *Dlx2* versus control vector (pcDNA3) co-transfection, after correction of the transfection efficiency with β-galactosidase activity. Error bars represent values of standard deviation (SD); **** denotes *P*< 0.0001.

### Olig2 expression is increased in the ventral forebrain of *Dlx1/2* DKO mouse embryos

The above data support that DLX2 protein binds to the mouse *Olig2* gene locus *in vivo*, and validate the specificity of binding and its transcriptional repressive function *in vitro*. Next, we conducted experiments to determine whether DLX2 protein is required to repress Olig2 expression *in vivo*. Due to the partial functional redundancy of *Dlx1* and *Dlx2* (Anderson, Qiu, et al., 1997), we investigated *Olig2* expression in the forebrain of *Dlx1*/*2* double mutant mouse embryos. In coronal sections of *Dlx1/2* mutant telencephalon at E13.5, IF assays detected a remarkable increase in the number of OLIG2 expressing cells in the SVZ and MZ of MGE, when compared with the WT littermate control (**Figure 6A**, b, d). At the transcription level, our RT-qPCR assay results demonstrated that *Olig2* mRNA expression is significantly higher in the GE tissues of *Dlx1/2* DKO embryos at E13.5 (**Figure 6C**). These results confirm that there is expanded *Olig2* expression in *Dlx1/2* DKO embryonic forebrain, supporting the loss of transcriptional repression of *Olig2* by DLX1 and/or DLX2 in the absence of active *Dlx1*/*2* functions *in vivo*.

**Figure 6.**
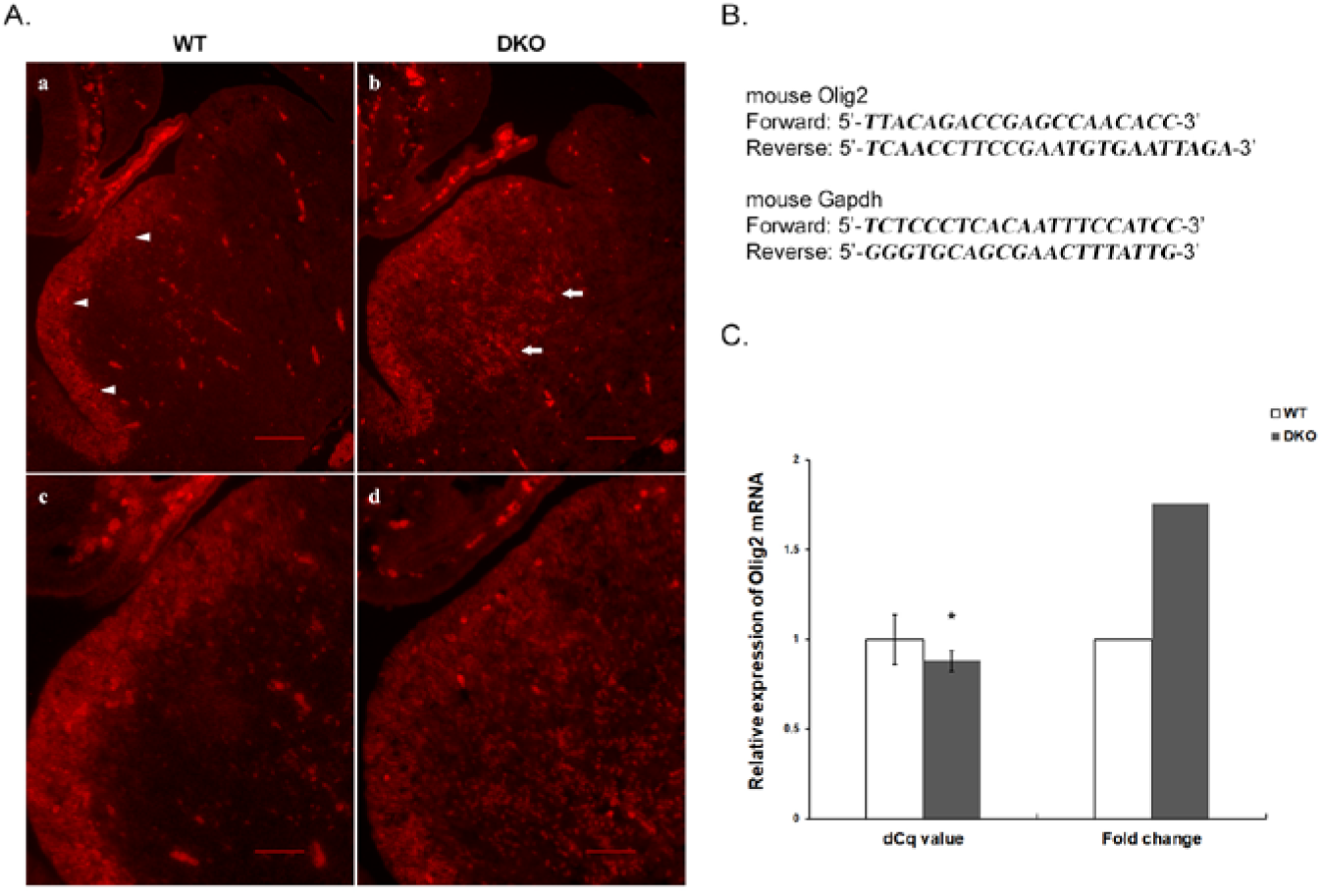
Olig2 expression is increased in E13.5 *Dlx1/2* DKO mouse forebrain. **A**, IF staining of OLIG2 on coronal sections of E13.5 WT (a, c) and *Dlx1/2* DKO (b, d) mouse embryonic forebrain, **c** and **d** are higher-magnification views of **a** and **b**, respectively. In WT control (**a**), arrowheads point to the region where OLIG2 expression decreases sharply in the SVZ of MGE coincidental with the presence of *Dlx1 and Dlx2*. This region is much less evident in *Dlx1/2* DKO mouse forebrain (**b**) in that the number of cells expressing OLIG2 is increased in the SVZ and MZ (arrows). **B**, The sequence of oligonucleotide primers used for qPCR is listed for mouse *Olig2* and *Gapdh* genes. **C**, RT-qPCR demonstrates higher *Olig2* mRNA expression in the GEs of *Dlx1/2* DKO embryos compared with WT littermates at E13.5. The dCq value indicates comparison of the ΔCq values of *Olig2* after normalization between *Dlx1/2* DKO and WT embryos. Error bars represent values of SD, and * denotes *P* < 0.05. The fold change is calculated using 2 ^ (-ΔΔCq) method. Scale bars: **A, a, b**, 100 µm; **c, d**, 50 µm.

### *Dlx1* represses *Olig2* expression in embryonic chicken forebrain

To further confirm the negative regulation of *Dlx* homeobox genes on *Olig2* expression, we performed *in ovo* electroporation to overexpress chicken *Dlx1* in *Olig2* expressing cells in the developing chicken forebrain. The GFP expressing RCASBP(B) retrovirus vector was used to deliver chicken *Dlx1*, or Q50E mutant *Dlx1* to the forebrain of E2 chicken embryos unilaterally. Electroporated embryos were harvested on E5, and *Dlx1* overexpression was confirmed by ISH on the coronal forebrain sections of the embryos (**Figure 7A**, a, e). IF assays on the adjacent sections showed that in GFP expressing cells (**Figure 7A**, b, f), which correspond to cells with *cDlx1* overexpression, the expression of OLIG2 was dramatically reduced or ablated (**Figure 7A**, c, g), without apparent co-expression of GFP and OLIG2 (**Figure 7A**, d, h). In control embryos electroporated with RCASBP(B) vector alone, ISH demonstrated endogenous *cDlx1* expression (**Figure 7B**, a, e) while no effect on OLIG2 expression was observed (**Figure 7B**, c, g), with numerous cells co-expressing both GFP and OLIG2 (**Figure 7B**, d, h). In embryos electroporated with Q50E mutant *cDlx1*, ISH produced robust overexpression signal because there is only one nucleotide change between the wild type and the mutant *cDlx1* (Figure 7C, a, e). However, this Q50E mutation at the homeodomain amino acid position 50 has been shown to be sufficient to eliminate the DNA binding ability of mouse DLX1/2 proteins *in vitro* (Le et al., 2007). Our EMSA findings confirmed that chicken DLX1 with a Q50E mutation is unable to bind the *Olig2* probe (**Figure 4C**, d). Our electroporation data further demonstrated that overexpression of Q50E mutant *cDlx1* does not have an inhibitory effect on OLIG2 expression in the embryonic chicken forebrain (**Figure 7C**, c, d, g, h). Together, our data indicate that chicken *Dlx1* is sufficient to repress OLIG2 expression cell-autonomously *in ovo*. Moreover, the Q50E mutation within the chicken DLX1 homeodomain leading to loss of its DNA binding capability results in abrogation of this repression, supporting that the observed repressive effect of DLX1 on *Olig2* expression is also direct transcriptional in the embryonic chicken forebrain.

**Figure 7.**
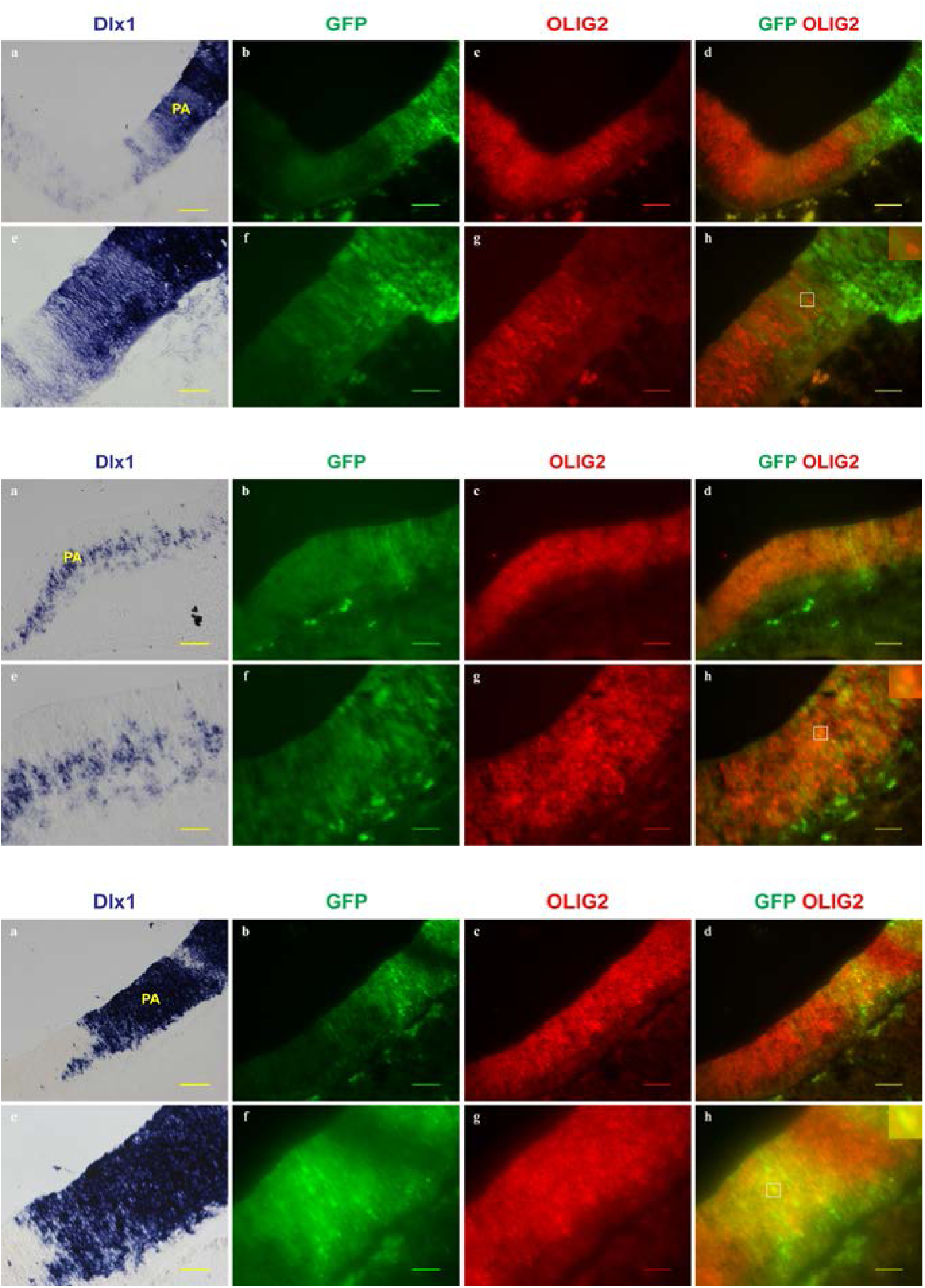
Over-expression of *Dlx1* in chicken telencephalon reduces OLIG2 expression *in ovo*. **A-C**, Images of coronal forebrain sections of E5 electroporated chicken embryos. ISH was performed to examine *cDlx1* expression status, while OLIG2 expression (red) in GFP positive electroporated cells was visualized using IF staining on the adjacent sections. All images are oriented with the electroporated side to the right and dorsal to the top. (**A**) In embryos electroporated with RCASBP(B)-*Dlx1*, there is intense ISH signal of *Dlx1* overexpression in the targeted side of the ventral forebrain (**a, e**). In GFP positive cells within the corresponding region on the adjacent section (**b, f**), OLIG2 staining declines (**c, g**) with minimal co-localization of GFP and OLIG2 (**d, h**). (**B**) In control embryos electroporated with RCASBP(B), the expression of *Dlx1* (**a, e**) and OLIG2 (**c, g**) does not change in GFP expressing cells (**b, f**). (**C**) Over-expression of mutant chicken *Dlx1* in embryos electroporated with RCASBP(B)-*Dlx1* Q50E (**a, e**); OLIG2 expression remains unaffected (**c, g**) with widespread overlap of GFP and OLIG2 staining (**d, h**), in a manner similar to embryos electroporated with RCASBP(B) alone (**B, d, h**). **A-C**, images **e-h** are higher magnification views of the PA domain of **a-d**, and insets in **h** highlight the relative localization of GFP and OLIG2 within the boxed region, respectively. PA, pallidum. Scale bars: **A-C, a-d**, 50 μm; **e-h**, 25 μm.

## DISCUSSION

Transcription factors interact in networks and play key roles at the core of the molecular and genomic mechanisms that regulate forebrain development and function (Nord, Pattabiraman, Visel, & Rubenstein, 2015). Precise regulation of the neuron-glial cell fate decisions among common progenitors is critical for modulating the balance of interneurons versus oligodendrocytes production, and therefore the normal development of the telencephalon. Previous work has established that *Dlx1/2* may promote interneuron differentiation while concurrently inhibiting oligodendrocyte specification through repression of *Olig2* expression in common bipotent progenitors within the ventral forebrain (Petryniak et al., 2007). In this study, we provide novel evidence that the effect of DLX on *Olig2* expression is direct transcriptional repression mediated by binding to the *Olig2* gene locus.

Four of the six known mouse Dlx genes (Dlx1, 2, 5 and 6) are expressed in the developing forebrain with both temporal and spatial overlap, suggesting potential redundant functions among these genes (Eisenstat et al., 1999; Liu et al., 1997). Indeed, while mice with single mutation of either *Dlx1* or *Dlx2* exhibit distinct but subtle defects in forebrain development (Cobos et al., 2005; Qiu et al., 1995), *Dlx1/2* double mutants have a severe deficiency in subcortical neurogenesis, an almost complete loss of tangential interneuron migration, as well as an increase in oligodendrogenesis (Anderson, Eisenstat, et al., 1997; Anderson, Qiu, et al., 1997; Long et al., 2007; Marin et al., 2000; Petryniak et al., 2007; Pleasure et al., 2000). *Dlx1* is proposed to be the founding member of the vertebrate *Dlx* family because it is most closely related to the *Dll* gene of Drosophila and amphioxus (Panganiban & Rubenstein, 2002). However, DLX2 is expressed before DLX1 during forebrain development in the mouse (Eisenstat et al., 1999; Le et al., 2017), and previously we also found that DLX2 is more robust than DLX1 as a transcriptional activator (Le et al., 2017; Q. P. Zhou et al., 2004) or transcriptional repressor (Le et al., 2007), signifying a more important role for the *Dlx2* gene in forebrain development.

We therefore focused our study on *Dlx2*. Interestingly, *Dlx2*’s sequence information was not available in the chicken genome assembly Gallus_gallus-4.0 when we initially tried to clone chicken *Dlx2*. Furthermore, when blasting the Gallus_gallus-4.0 database with mouse *Dlx2* ORF sequence, no other potential *Dlx* sequence was found on chicken chromosome 7, except *Dlx1*, while *Dlx1* and *Dlx2* are known to be arranged in one bigenic cluster on the same chromosome in both human and mouse (Panganiban & Rubenstein, 2002). Although there was a predicted chicken *Dlx2* sequence in the updated chicken genome assembly (Gallus_gallus-5.0), two pairs of PCR primers based on the predicted sequence failed to amplify the potential chicken *Dlx2* using the same E7 chicken retina cDNA template for *Dlx1*. According to our previous findings, both *Dlx1* and *Dlx2* are expressed in developing mouse retina by E12.5 (Eisenstat et al., 1999). Dlx2 expression in the mouse retina is maintained throughout the lifetime while *Dlx1* expression decreases postnatally (de Melo et al., 2003). Thus, it is highly possible that chicken has lost the *Dlx2* ortholog during evolution. Collectively, our data about chicken *Dlx1* and mouse *Dlx2* also suggest that DLX1 or DLX2 alone is sufficient to repress the expression of *Olig2*. However, because of the cross-regulatory interactions between members of the *Dlx* gene family, mutation of *Dlx1/2* leads to loss of *Dlx5/6* expression in the SVZ of the developing striatum (Anderson, Qiu, et al., 1997; Zerucha et al., 2000). It is unclear whether this loss of *Dlx5/6* expression contributes partially to the increase of oligodendrogenesis in the *Dlx1/2* mutants, as the regulatory roles that *Dlx5* and/or *Dlx6* play in the expression of *Olig2* have not yet been determined (B. Wang, Lufkin, & Rubenstein, 2011; Y. Wang et al., 2010).

Our expression pattern analysis of *Dlx1/2* and *Olig2* in chicken and mouse provides evidence to support the model of interneuron versus oligodendrocyte specification in the basal telencephalon as described (Petryniak et al., 2007). In general, *Olig2* defines one bipotent progenitor domain in the VZ. *Dlx1* in chicken and *Dlx2* in mouse transiently co-express with *Olig2* in a subset of these progenitors. After cells exit the progenitor zone and begin to differentiate, the expression of *Dlx1/2* and *Olig2* becomes mutually exclusive with observation of an increase of *Dlx1/2* with concurrent reduction of *Olig2* expression (**Figure 1**, **2**). Additional evidence of this *Dlx* mediated repression of Olig2 expression is obtained from the *Dlx1/2* mouse mutants. In the absence of functional *Dlx1/2* inhibition, *Olig2* expression is significantly increased in *Dlx1/2* DKO embryonic forebrain (**Figure 6**). Our overexpression data in chicken indicate that *cDlx1* is sufficient to repress OLIG2 expression cell-autonomously *in ovo* (**Figure 7**). Our findings further establish that *Dlx1/2* regulation of Olig2 expression is via transcriptional repression mediated by direct binding. First, we validated the *in vivo* binding of DLX2 protein to the *Olig2* gene locus in the mouse embryonic forebrain (**Figure 3**). ChIP is the most widely used technique to identify protein-DNA associations *in vivo* (Fullwood & Ruan, 2009). We have optimized our ChIP protocol to preferentially obtain homeoprotein-genomic DNA complexes from embryonic tissues *in situ* (Q. P. Zhou et al., 2004). With the sensitive and highly specific antisera we possess (Eisenstat et al., 1999), several direct transcriptional targets of DLX1/2 proteins during forebrain development have been determined including the *Dlx5/6* intergenic enhancer, *neuropilin-2* and the glutamic acid decarboxylase genes *Gad1/Gad2* (Le et al., 2007; Le et al., 2017; Q. P. Zhou et al., 2004). Second, our EMSA data confirmed that mouse DLX2 protein is capable of binding to mR2 and mR6 regions of mouse Olig2 gene *in vitro* (**Figure 4** A, B). Third, we found that DLX2 represses transcription of a reporter vector containing the mR6 sequence of the mouse *Olig2* gene *in vitro* (**Figure 5**). On the other hand, due to absence of an antibody for chicken DLX1, we have not obtained data of chicken DLX1 binding to the *Olig2* locus *in ovo*. However, our EMSA results demonstrated the binding of chicken DLX1 to the cR1 domain of the chicken *Olig2* gene *in vitro*, a region showing a high degree of sequence similarity with the mouse Olig2 mR2 region *in silico* (**Figure 4** C). Given that there is no homeodomain binding element in the chicken *Olig2* locus that corresponds to the mR6 region of the mouse *Olig2* gene, we propose that mouse DLX2 may directly repress *Olig2* transcription through two mechanisms via binding to the mR2 and mR6 regions, respectively, while chicken DLX1 functions via the cR1 sequence sharing the same repression mechanism with the mouse *Olig2* mR2 domain. In chicken, the Q50E homeodomain mutant of DLX1 was unable to bind to the *Olig2* probe *in vitro*, and loses its ability to repress OLIG2 expression *in ovo*, strongly supporting that this observed repressive effect by cDLX1 on *Olig2* expression is also directly transcriptional.

The detailed mechanisms by which DLX1/2 repress *Olig2* expression remain unclear. It has been reported that the LIM domain-binding protein 1 (LDB1) is capable of mediating promoter:enhancer looping through LDB1 homodimerization (Deng et al., 2012; Krivega, Dale, & Dean, 2014). In a pituitary cell type, LDB1 can mediate transcriptional repression via promoter:enhancer looping by recruiting negative cargo to repressive enhancers, which results in promoter pausing of the targeted genes. Although LDB1 is bound predominantly to enhancers, enhancer:promoter-dependent LDB1 homodimerization may not account for the majority of LDB1-mediated looping events (Zhang et al., 2015). Our findings in mouse demonstrate binding of DLX2 protein to the *Olig2* locus at mR6i and to the canonical homeodomain TAAT binding motif within the mR2 region about 1.9 kb upstream (**Figure 3** A). This raises the possibility of DNA looping mediated transcriptional repression of *Olig2* by DLX2. However, since the chicken has only the cR1 domain that corresponds to the mouse mR2 region, without a comparable binding motif in mouse mR6i (**Figure** 4 C, b), this possibility is less likely. Notably, the mouse mR6i motif is located in the noncoding exon 1 of the mouse *Olig2* gene.

It has been reported that the ubiquitous transcriptional repressor CCAAT-displacement protein CDP/Cux binds to the repressor element in untranslated exon 1 and represses the transcription of the bi-exonic *Rnf35* gene in early mouse development (Huang, Chang, Wu, & Choo, 2005). The *Rnf35* gene uses an initiator overlapping with the 5’ end sequence of the exon 1 as the sole core promoter element in transcription, and requires nuclear factor Y (NF-Y) targeting a Y-box element immediately upstream of the initiator as a transcriptional activator (Huang, Wu, & Choo, 2005), while the CR1-CR2 segment of CDP has been shown to be able to displace NF-Y binding *in vitro* (Moon, Berube, & Nepveu, 2000). The underlying mechanism of the *Olig2* mR6i motif modulated transcriptional repression by DLX2 still requires active investigation. As for the cR1 *Olig2* motif of chicken and the corresponding mR2 *Olig2* domain of mouse, we postulate that possible associated mechanisms of transcriptional repression by DLX1/2 may include long-range repression via chromatin remodelling mediated with co-repressors, and/or ablation of activator function (Courey & Jia, 2001; Gaston & Jayaraman, 2003), as the mR2 domain alone did not show a repressive effect in luciferase reporter assays using HEK293 cells *in vitro* (**Figure 5**).

Interestingly, it has been reported that most of the homeodomain transcription factors expressed by neural progenitor cells in the ventral neural tube share a conserved domain related to core repression motif eh1 of the engrailed homeoprotein. These homeodomain proteins recruit the Groucho/TLE/Grg co-repressors with the eh1-like domain, and function as transcriptional repressors to establish neural patterning *in vivo* (Muhr, Andersson, Persson, Jessell, & Ericson, 2001). Further evidence suggests that histone deacetylase (HDAC) is required for the repression of *Olig2* by NKX2.2 in the developing chicken neural tube (H. Wang & Matise, 2016), while HDAC has been identified as one interacting protein, and thus can repress transcription together with the bridging Groucho/TLE/Grg protein in a corepressor complex by inducing a non-permissive chromatin structure (Winkler, Ponce, & Courey, 2010).

